# The impact of bone cancer on the peripheral encoding of mechanical pressure stimuli

**DOI:** 10.1101/498980

**Authors:** Mateusz W. Kucharczyk, Kim I. Chisholm, Franziska Denk, Anthony H. Dickenson, Kirsty Bannister, Stephen B. McMahon

**Author notes:** Correspondence, Central Modulation of Pain Group, Wolfson Centre for Age-Related Diseases, King’s College London, London SE1 1UL, UK.

## Abstract

Skeletal metastases are frequently accompanied by chronic pain that is mechanoceptive in nature and not easily managed by available therapies. The peripheral sensory profile of primary afferents responsible for transmitting the pain-related messages from cancerous bone to central sites is investigated here. We imaged thousands of primary sensory dorsal root ganglion neurons *in vivo* in healthy (sham-operated) and cancer-induced bone pain (CIBP) rats in order to analyse and compare their function. Utilising Markov Cluster Analysis we identified distinct clusters of primary afferent responses to limb compression and position. In CIBP rats, three times as many sensory afferents responded to knee compression in the leg ipsilateral to the tumour compared to sham-operated rats. We present evidence that the observed increase in sensory afferent response was not due to increased individual afferent activity but rather represents activation of ‘silent’ nociceptors, whose origin we propose is largely from outside of the bone.

## Introduction

A significant side effect of certain cancer types is severe pain, which can persist even in remission (Clohisy & Mantyh, 2003; Mantyh, 2006; Mantyh, Clohisy, Koltzenburg, & Hunt, 2002). Cancer pain is particularly prominent in cases where malignant tumours have invaded the skeleton (Mantyh, 2006; Nyquist & Nelson, 2017). This type of mechanoceptive pain, is difficult to manage in mobile subjects partly because it has considerable ‘break-through’ pain components (Mantyh et al., 2002).

Mechanistically, cancer-induced bone pain (CIBP) is transmitted via peripheral pseudounipolar somatosensory neurons that are genetically and physiologically heterogenous (Chisholm, Khovanov, Lopes, Russa, & McMahon, 2018; Lopes, Denk, & McMahon, 2017; Prescott, Ma, & De Koninck, 2014a; Schmelz, Schmid, Handwerker, & Torebjörk, 2000; Usoskin et al., 2014; Wang et al., 2018; Zeisel et al., 2018). How exactly they encode stimulus intensity on the populational level remains only partly explained (Prescott et al., 2014a; Wang et al., 2018). Two mechanisms are hypothesised: frequency coding (a number of action potentials fired in response to a given stimulus strength) and populational coding (the number of afferents engaged in response to a given stimulus). *In vivo* imaging of genetically-encoded calcium indicators offers the ability to sample large neuronal populations. Utilising this technique others reported that heating and cooling sensations are encoded differently by primary afferents (Wang et al., 2018). Despite pioneering work, many questions remain regarding afferent function. In this study we used *in vivo* functional imaging of rat somatosensory neurons to investigate how they encode mechanical stimuli during compression and by altering limb position and whether these modalities are encoded by musculoskeletal afferents. Crucially we compare functional responses in healthy control and CIBP rats using an unsupervised clustering of neuronal responses to reveal major responder classes to defined stimuli.

## Results and Discussion

### The impact of cancer progression on bone innervation

Pain associated with bone cancer is hypothesised to arise from altered primary sensory functionality. We first surgically delivered syngeneic mammary gland carcinoma (MRMT1) to a rat tibia (Medhurst et al., 2002) and evaluated the extent of bone damage (caused by tumour growth) using a micro-computer tomography technique (μCT) at two different time points: days 7/8 and days 14/15. Significant damage to the trabecular bone was observed at both stages, but only cortical bone integrity was compromised at the later time stage (days 14/15). As such our model has distinct early and late stage cancer components (Fig. 1A, Movie 1). Volumetric reconstruction enabled bone mineral density (BMD) quantification for the whole tumour growth area (Fig. S1A). Reduction of trabecular BMD was observed in both sham-operated and CIBP early and late stage rat groups (Fig. 1B), but cortical BMD was reduced only in the late stage CIBP rats (485 mg/cm^3^ in the sham-operated rat group compared to 146 mg/cm^3^ in the CIBP late stage group) (Fig. 1C). This was reflected in visually evident bone lesions.

**Figure 1.**
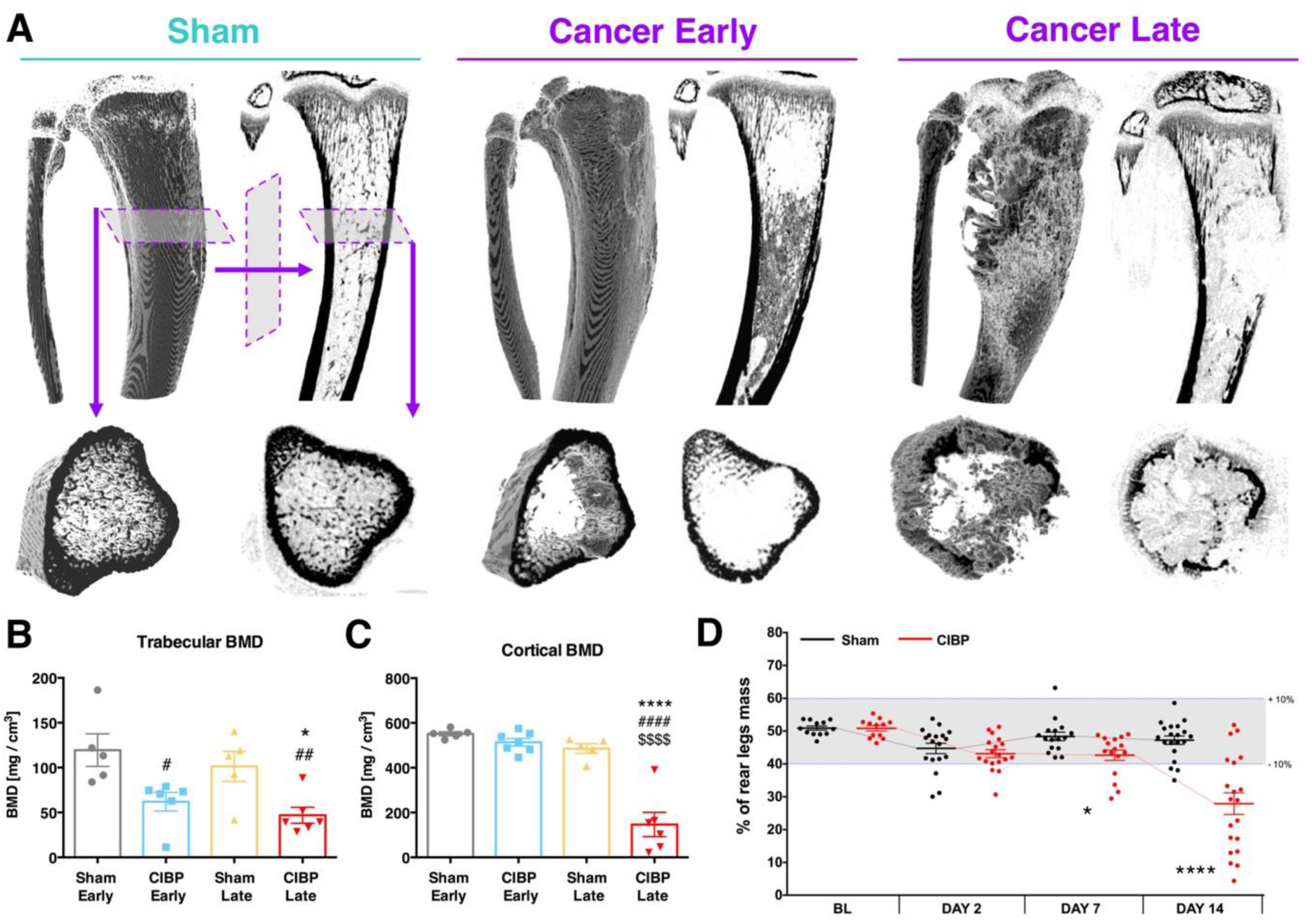
The impact of cancer progression on bone innervation. Example micro-computer tomography reconstructions of rat tibiae. Panels depict a sham-operated control, an early (day 7/8) and a late (day 14/15) cancer stage. Top panels represent a whole 3D-rendered tibia with corresponding orthogonal projection. Bottom panel shows a plantar representation of selected micro-scans from the top panel. An early cancer stage is characterised by the trabecular bone lesions whereas the late stage by both trabecular and cortical bone lesions **(A)**. Trabecular bone mineral density quantification. Volumetric bone mineral density quantification from 114 reconstructed micro-scans (every 34 *µ*m) per bone. Selected planes for analysis were chosen to cover the tumour growth area (see methods for details). Each dot represents a single bone from a separate animal (n = 5-7 per group). Data represent the mean ± SEM (in mg/cm^3^). One-Way ANOVA: F_3, 18_ = 6.272, p = 0.0042 (group), with Tukey post-hoc test: # vs. Early Sham, * vs. respective Sham. * or ^#^ p < 0.05, ^##^ p < 0.01 **(B).** Cortical bone mineral density quantification. Volumetric bone mineral density quantification from 114 reconstructed micro-scans (every 34 *µ*m) per bone. Selected planes for analysis were chosen to cover the tumour growth area (see methods for details). Each dot represents a single bone from a separate animal (n = 5-7 per group). Data represent the mean ± SEM (in mg/cm^3^). One-Way ANOVA: F_3, 19_ = 35.57, p < 0.0001 (group), with Tukey post-hoc test: # vs. Early Sham, * vs. respective Sham, $ vs. Early Cancer, ****p < 0.0001 **(C)**. Static weight bearing measurement of rear legs. Within a timepoint, each dot represents a single animal (n = 13-20 per group). Each measurement was taken as an average of 5 consecutive readouts per animal per timepoint. Data represent the mean ± SEM. Kruskal-Wallis H for independent samples (Cancer vs. Sham): day 0: χ^2^(1) = 0.083, p = 0.773, day 2: χ^2^(1) = 1.602, p = 0.206, day 7: χ^2^(1) = 5.286, p = 0.022, day 14: χ^2^(1) = 15.384, p < 0.0001. *p < 0.05, ****p < 0.0001 vs. respective Sham **(D)**. See also Figure S1 and Movie S1.

Tumour progression is expected to relate to animal behaviour, and we monitored rats up to 14 days after surgery. While body weight gain remained stable in all groups (Fig. S1B), the behavioural data demonstrate that CIBP rats manifest mechanical hypersensitivity; significant changes in static weight bearing between rear legs is evident from day 7 post-surgery (Fig. 1D). These results correspond to other studies using similar rodent models of CIBP (Medhurst et al., 2002; Urch, Donovan-Rodriguez, & Dickenson, 2003).

### The number of mechanically-responsive sensory neurons is tripled in animals with bone cancer

Because cortical BMD was reduced only in late stage CIBP rats, all data considered forthwith is from late stage sham-operated and CIBP rats. We studied mechanosensation using *in vivo* imaging with GCaMP6s following intrathecal delivery of an AAV9 vector containing GCaMP6s (timeline depicted Fig. 2A, imaging and stimulation sites depicted Fig. 2E). This method ensures uniformly distributed expression between all subtypes of DRG neurons (Chisholm et al., 2018) (Fig. S1C). In total we analysed 757 DRG neuronal cell bodies from 9 sham-operated rats, and 1748 neurons from 10 CIBP animals (Fig. 2B, C). Lumbar DRG L3 and L4 were chosen for imaging based on our fast blue (FB) tracing studies (Fig. S1C, and (Kaan et al., 2010; Peters et al., 2005)), and 6-7 L3 DRG and 3 L4 DRG from each group were imaged. No response difference could be detected between L3 and L4 lumbar levels (not shown) and so results from all DRG were pooled.

**Figure 2.**
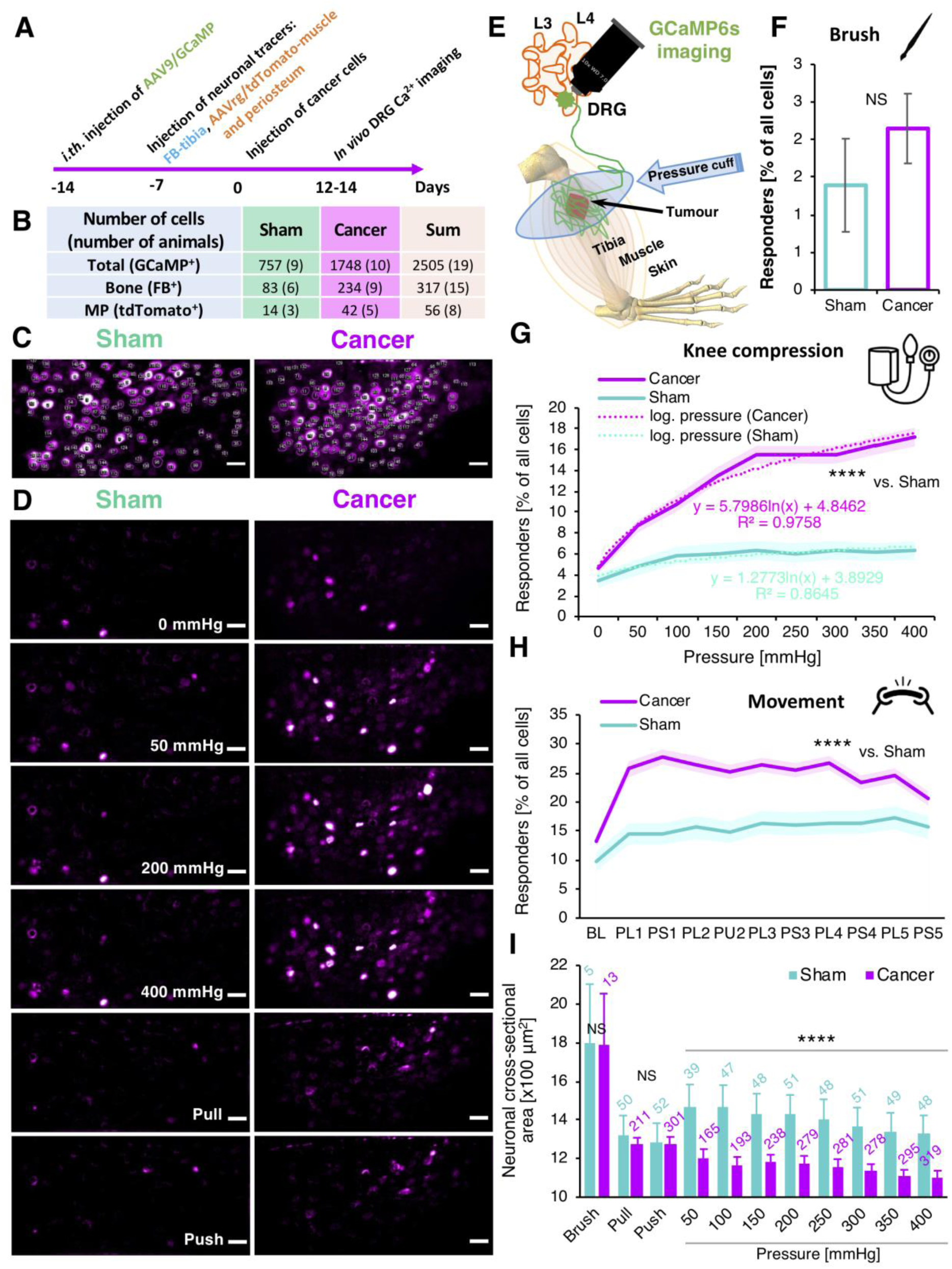
The number of mechanically-responsive sensory neurons is tripled in animals with bone cancer. An experimental timeline. The injection of neuronal tracers and cancer cells implantation was performed via the same hole in the tibia to limit damage. FB – fast blue, GCaMP6s – genetically encoded calcium indicator, AAV9 – adeno-associated virus serotype 9, AAVrg – adeno-associated virus serotype 2 retrograde **(A)**. A summary table representing the number of cells analysed in the *in vivo* calcium imaging experiment. In brackets are the numbers of animals studied **(B)**. Representative frames from the end of an *in vivo* imaging session with circled DRG neuronal cell bodies for analysis. The confocal z-scan was taken over 30 minutes *post mortem* allowing for calcium to build up in the neuronal cell bodies permitting indication of most of the GCaMP6s labelled cells **(C)**. Selected video frames of imaged DRG cell bodies responding to the knee compression and the leg movements (Push/Pull along the body axis). Sham-operated (left) and Cancer rat model (right). Note the increase in number of responding neurons between the groups. Scale bars, 100 *µ*m **(D)**. Schematic representation of the *in vivo* GCaMP6s imaging. DRG, dorsal root ganglia; GCaMP6s, genetically encoded calcium indicator, L3, L4 are lumbar vertebrae 3 and 4, respectively. Pressure was delivered utilising neonatal pressure cuff surrounding the animal’s knee. Rats were stably anaesthetised with urethane **(E)**. Percentage of responders from 1337 neuronal cell bodies analysed from L3 and L4 DRG during dynamic brushing of the leg surface in Sham and Cancer animals. The denominator for the total number of cells was established at the end of the experiment as explained in (C). Data represent the mean ± SEM (shaded areas) of n = 359 cells (sham) from 6 animals, and n = 978 cells from 6 animals (Cancer). Student t-test: p = 0.376 **(F)**. Percentage of responders from 2505 neuronal cell bodies analysed from L3 and L4 DRG during knee compression in Sham and Cancer animals. Pressure in the neonatal cuff overlaying knee joint was increased in 50 mmHg steps every 10 s (see methods for more detail). The denominator for the total number of cells was established at the end of the experiment as shown in (C). Data represent the mean ± SEM (shaded areas) of n = 757 cells (sham) from 9 animals, and n = 1748 cells from 10 animals (Cancer). The increase in number of responding cells follows the logarithmic function of the pressure applied. RM-ANOVA: F_1, 2503_ = 43.276, p < 0.0001 (vs. sham) **(G)**. Percentage of responders from 1592 neuronal cell bodies analysed from L3 and L4 DRG during the gentle leg movement along the body axis in Sham and Cancer animals. Responses of 5 consecutive pull (PU) and push (PS) pairs are presented. The denominator for the total number of cells was established at the end of the experiment as shown in (C). Data represent the mean ± SEM (shaded areas) of n = 359 cells (sham) from 6 animals, and n = 1233 cells from 7 animals (Cancer). RM-ANOVA: F_1, 1590_ = 17.396, p < 0.0001 (vs. sham) **(H)**. Cell size analysis of all responders during: dynamic brushing of the leg surface, leg movement along the body axis (first pull and push shown), and knee compression in Sham and Cancer animals. Pressure was increased in 50 mmHg steps every 10 s. Data represent the mean ± SEM (shaded areas). Brush: unpaired t-test: p = 0.984, Movement: Kruskal-Wallis for independent samples: F_1, 674_ = 0.110, p = 0.740, Knee compression: Kruskal-Wallis for independent samples: F_1, 2429_ = 28.469, p < 0.0001. Number of cells analysed are in brackets above each bar **(I)**. See also Figure S2 and Movie S2.

The ipsilateral hind limb was stimulated in three ways: brush (Fig. 2F), movement of the limb along the body axis in 5 consecutive push-pull stretching cycles (Fig. 2H), and leg compression delivered by a neonatal pressure cuff with separate overlay for the knee and tibial head, calf, and lower calf-ankle; an incremental pressure (50 mmHg every 10 s, in the range of 0-400 mmHg) was applied (Fig. 2G, S2A, B). No significant difference in the number of primary afferent responders to dynamic brushing between sham-operated and CIBP animals was observed (Fig. 2F). In contrast, quantitative analysis of the percentage of responding cells in sham-operated versus CIBP rats following knee (the tumour-growth area) compression indicated a 3-fold increase in the number of responders in CIBP rats (to around 18% of L3/4 DRG neurones) rats compared to their sham-operated counterparts (Fig. 2G, S2A, B). Visual representations are provided for before and after stimulation (Movie 2, Fig. 2D). Interestingly, increased compression force was reflected in the recruitment of responders in the CIBP group in a logarithmic fashion (Fig. 2G). CIBP rats also demonstrated a 2-fold increase in primary afferent responders following limb movement (without load bearing) compared to sham-operated rats (Fig. 2H).

We analysed responder cell size distribution in sham-operated and CIBP rats (choosing 700 *µ*m^2^ and 1200 *µ*m^2^ to separate small and medium-size cells). Our results suggest that pressure is encoded mainly by small to medium-size neurons. In CIBP rats a decrease in average responding cell size with increased compression force is suggestive of small nociceptive afferent recruitment (Fig. S2D). We hypothesise that these were previously ‘silent’ nociceptive afferents, the recruitment of which has been reported previously in inflammatory conditions (Feng et al., 2012; Gold & Gebhart, 2010; Prato et al., 2017; Schaible & Schmidt, 1985). This type of afferents particularly innervates deep body structures (up to 50% of all deep afferents are ‘silent’ in mice (Feng et al., 2012; Prato et al., 2017; Schaible & Schmidt, 1985)), but rarely rodent’s skin (Wetzel et al., 2007). Recently, a potential mechanism for ‘unsilencing’ of silent nociceptors by NGF-TrkA-Piezo2 has been proposed (Prato et al., 2017). Considering the observed increase in the number of responders in cancer conditions as well as the fact that the CIBP is NGF-dependent (Bloom et al., 2011; Tomotsuka et al., 2014), it is reasonable to predict that silent nociceptors would significantly contribute to the development of pain in the CIBP rats. In sham-operated rats the number of medium sized responders increased with increasing stimulus pressure while dynamic brushing only recruited few large-sized neurons. This correlates with previous literature regarding the preferential encoding of noxious compression by myelinated Aδ nociceptors (Nencini & Ivanusic, 2017), while light touch is encoded by large Aβ fibres (Chisholm et al., 2018; Ma & Woolf, 1996) (Fig. 2I, S2).

### Leg compression and position are differentially coded by DRG sensory neurons

Contrary to cell number, the fluorescence intensity of responders was not altered, on average, in CIBP rats, with no difference observed between groups after mechanical stimulation. However, fluorescence intensity increased proportionally with pressure intensity in both sham-operated and CIBP rats (Fig. 3A, B, C, S4A, B). The major responder pattern of firing was analysed using Markov Cluster Analysis (MCA). This largely unsupervised approach revealed 4 main clusters of neuronal responses to limb compression where afferent responses were classified as (1) triggered by ‘low’ pressure (<100 mmHg), (2) triggered by ‘middle’ pressure (peak at around 200 mmHg), (3) triggered by ‘high’ pressure (>300 mmHg), or (3) triggered by the pressure surge (0-400 mmHg, ‘ramp’) (Fig. 3D). High but not low responders likely represent nociceptive-specific afferents. The most abundant ramp cluster likely reflects graded frequency coding of compressive forces (similarly to heat (Wang et al., 2018)), which is maintained in both health and disease (Prescott, Ma, & De Koninck, 2014b). Neuronal responses to limb movement revealed 3 clusters, classified as a fluorescence increase to limb (1) ‘pull’, (2) ‘push’ or (3) movement (‘const’) (Fig. 3E). The presumed proprioceptive response most likely reflects combinatorial coding, where different cells encode the different position of the limb (similarly to cold (Wang et al., 2018)) (Prescott et al., 2014b).

**Figure 3.**
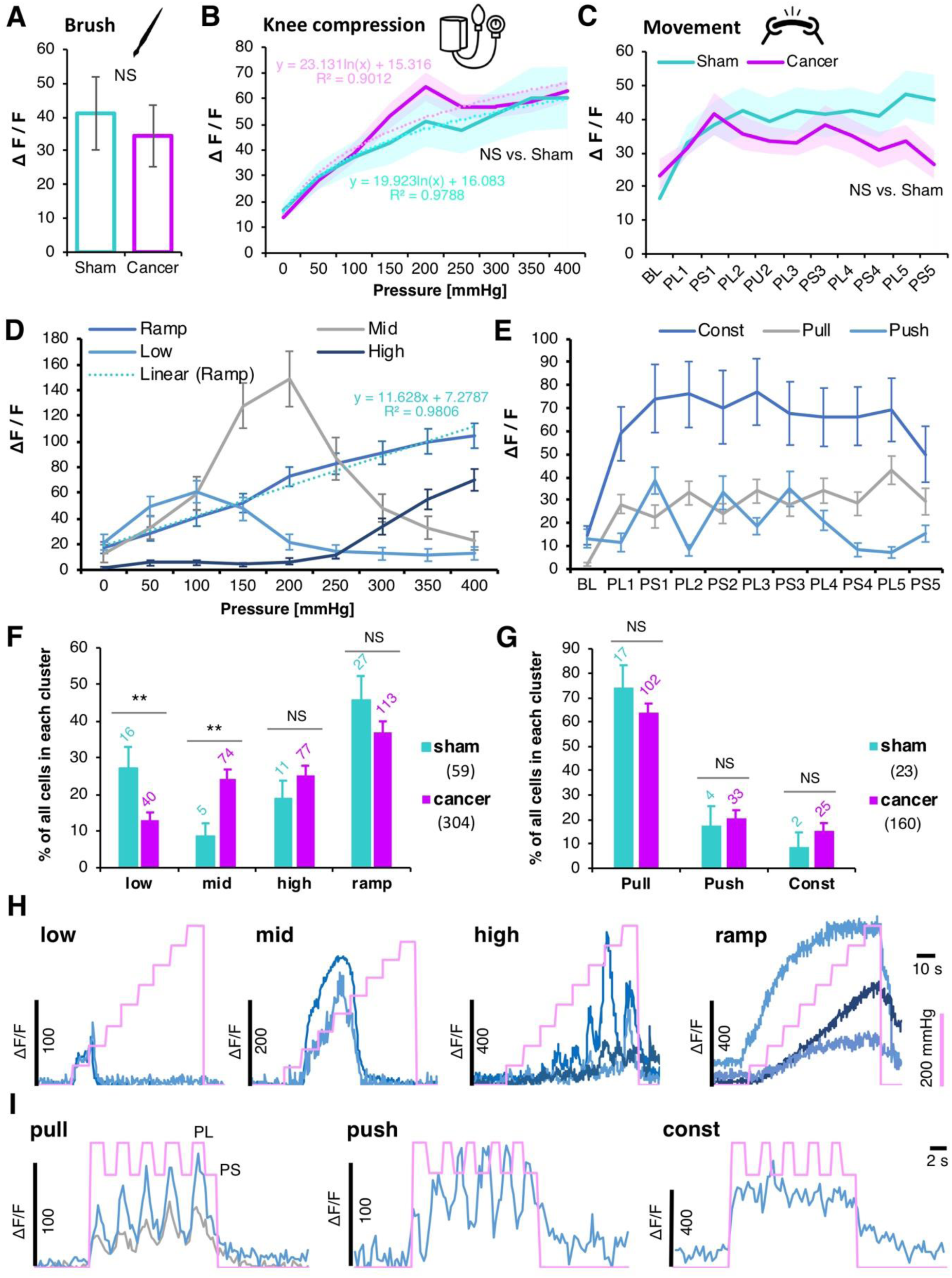
Leg compression and position are differentially coded by DRG sensory neurons. Normalised fluorescence intensities of all responding neuronal cell bodies from L3 and L4 DRG to dynamic brushing of the leg. Data represent the mean ± SEM. N = 5 cells (sham) from 6 animals, and n = 21 cells from 6 animals (Cancer). Unpaired t-test: p = 0.737 **(A)**. Normalised fluorescence intensities of all responding neuronal cell bodies from L3 and L4 DRG to knee compression. Pressure was increased in 50 mmHg increments every 10 s. Data represent the mean ± SEM (shaded areas). N = 70 cells (sham) from 9 animals, and n = 415 cells from 10 animals (Cancer). RM-ANOVA: F_1, 483_ = 0.177, p = 0.674 (vs. sham) **(B)**. Normalised fluorescence intensities of all responding neuronal cell bodies from L3 and L4 DRG to leg movement along the body axis (PL – Pull, PS – Push). Data represent the mean ± SEM (shaded areas). N = 83 cells (sham) from 6 animals, and n = 493 cells from 7 animals (Cancer). RM-ANOVA: F_1, 574_ = 0.306, p = 0.580 (vs. sham) **(C)**. Unsupervised clustering of neuronal responses to the knee compression. Markov Clustering Analysis (MCA) revealed 4 major clusters of responders to knee compression; those that responded with the gradual increase of fluorescence in function of the pressure increase (Ramp), or those that responded only to the lower pressures (Low), middle pressures (Mid) and high pressures (High). Due to the lack of differences in the clusters between the Sham and Cancer animals, groups were pooled (see Figure S4) **(D)**. Clustering of neuronal responses to the limb’s movement. MCA revealed 3 major clusters of cells; those that responded stably throughout the movements (Const), or those that responded with the increase in fluorescence only to the leg pulling (Pull) or pushing (Push). Due to the lack of differences in the clusters between the Sham and Cancer animals, groups were pooled (see Figure S4) **(E)**. Percentage of cells in each knee compression cluster with regards to the group classifiers. Numbers over the bars represent total numbers of cells identified in each cluster. Total numbers of cells after MCA in Sham and Cancer groups are given in brackets. Note that ‘Mid’ cluster is particularly well represented in the cancer conditions. Unpaired t-test: ‘Low’ P = 0.007, ‘Mid’ P = 0.007, ‘High’ P = 0.274, ‘Ramp’ P = 0.216. **p < 0.01, NS – non significant **(F)**. Percentage of cells in each movement cluster with regards to the group classifiers. Numbers over the bars represent total numbers of cells identified in each cluster. Total numbers of cells after MCA in Sham and Cancer groups are given in brackets. Unpaired t-test: ‘Pull’ P = 0.342, ‘Push’ P = 0.720, ‘Const’ P = 0.384. NS – non significant **(G)**. Example normalised GCaMP6s fluorescence traces from DRG neurons of the identified knee compression clusters. Pink line indicates the surge in cuff pressure **(H)**. Example normalised GCaMP6s fluorescence traces from DRG neurons of the identified movement-evoked clusters. Pink line indicates events: higher points are reflecting pulling (PL) and lower pushing (PS) the leg along the body axis **(I)**. See also Figure S3, S4 and Movie S2.

There were significantly more cells in the mid, and less in the low, clusters in CIBP rats compared with sham-operated animals (Fig. 3F). This further suggests the recruitment of nociceptive, mechanically-responding cells in the disease. The decrease in low cluster cells could suggest an inhibition of Aβ-fibres, corresponding to the numbness experienced by patients, or the loss of this fibre type in CIBP rats. The lack of difference in the number of responders in the movement clusters suggests that no one limb position is particularly painful, simply that movement alone engages more cells in CIBP rather than sham-operated rats (Fig. 3G).

### Intratibial afferent function in health and bone cancer

As observed previously (Nencini & Ivanusic, 2017; Prato et al., 2017) we confirmed the presence of Piezo2 and TrkA proteins on bone afferents (Fig. 5A). Secondly, we investigated the functional responses of FB-traced intratibial afferents (Fig. 4A). Previously these fibres were shown to respond to intraosseous pressure in healthy animals (Nencini & Ivanusic, 2017; Nencini et al., 2017), while NGF sensitizes mechanically activated bone nociceptors (Nencini et al., 2017). We extend these findings by revealing that around 13% of traced tibial cavity afferents responded to knee compression with no differences observed between sham-operated and CIBP rats. This suggests that silent nociceptors in the late cancer stage originate from outside of the bone (Fig. 4B). High-pressure stimulation (around 200 mmHg) was required to engage FB responders, suggesting either that the bone has a higher-pressure activation threshold compared to extra-osseus afferents and/or that the same pressure values were not reached deep in the bone. Fluorescence intensity of bone afferents after the knee compression was equally unaffected by the presence of cancer (Fig. 4C).

**Figure 4.**
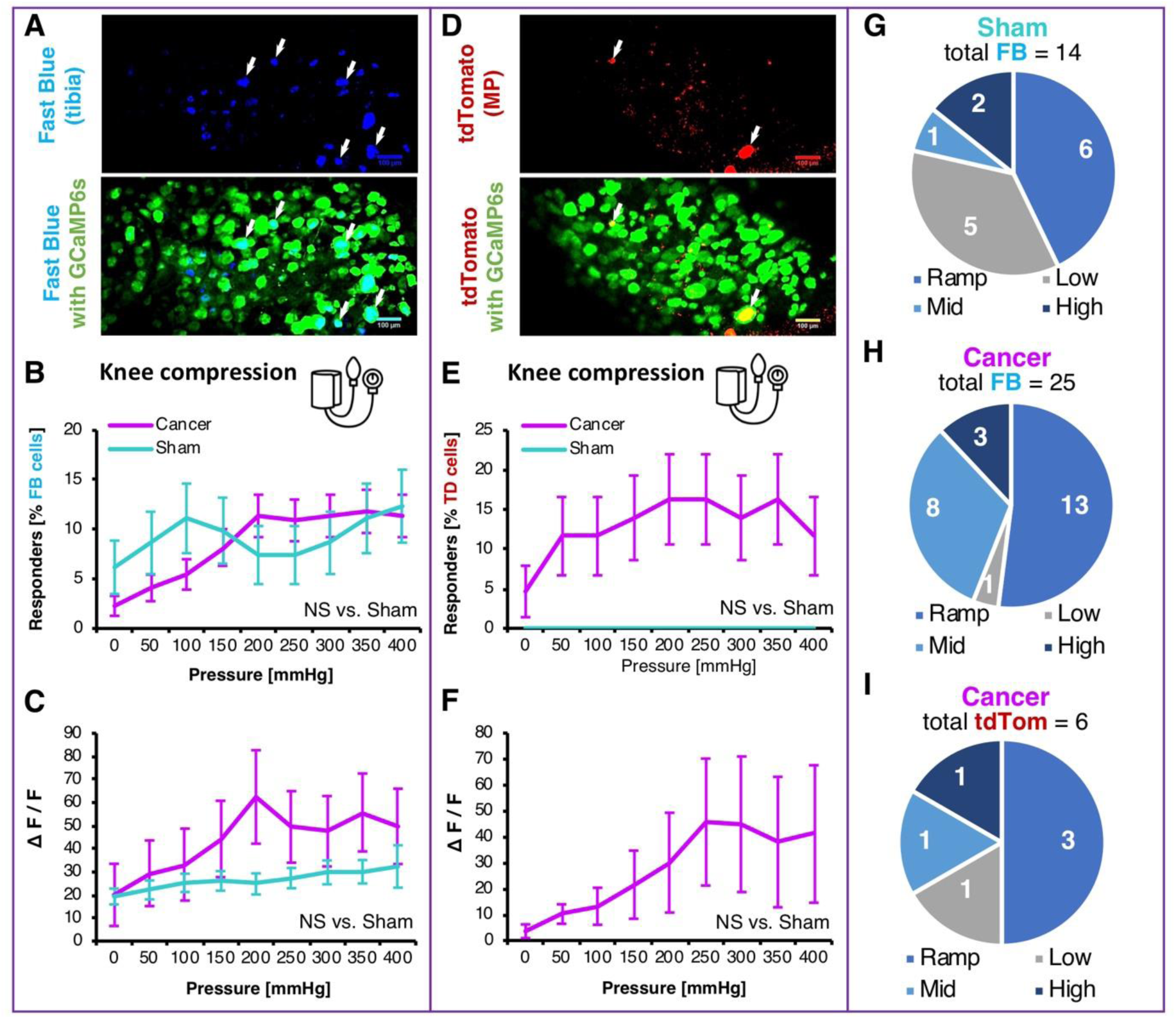
Intra- and peri-tibial afferent function in health and bone cancer. Fast Blue (FB) traced neurons from the tibial cavity. Representative z-stack collected at the end of the *in vivo* imaging experiment in order to identify traced cells (blue) and GCaMP6s (green). Scale bars: 100 *µ*m. Arrows indicate FB-traced neurons **(A)**. Percentage of responders of all FB-traced tibial afferents within L3 and L4 DRG during knee compression in Sham and Cancer animals. Pressure was increased in 50 mmHg steps every 10 s (See methods for more). Data represent the mean ± SEM of N = 81 cells from 6 animals (sham) and 220 cells from 9 animals (Cancer). RM-ANOVA: F_1, 299_ = 0.045, p = 0.832 **(B)**. Fluorescence intensities of all responding FB+ neuronal cell bodies to knee compression. Pressure was increased in 50 mmHg increments every 10 s. Data represent the mean ± SEM of N = 15 cells (Sham) from 6 animals, and n = 34 cells from 9 animals (CIBP). RM-ANOVA: F_1, 47_ = 0.552, p = 0.461 **(C)**. AAVrg/tdTomato (TD) traced neurons innervating the muscle and periosteum (MP) surrounding tibia. Representative z-stack collected at the end of the *in vivo* imaging experiment in order to identify traced cells (red) and GCaMP6s (green). Scale bars: 100 *µ*m. Arrows indicate TD-traced neurons **(D)**. Percentage of responders of all TD-traced MP afferents within L3 and L4 DRG during knee compression in Sham and Cancer animals. Pressure was increased in 50 mmHg steps every 10 s (See methods for more). Data represent the mean ± SEM of N = 14 cells from 3 animals (Sham) and 43 cells from 5 animals (Cancer). RM-ANOVA: F_1, 55_ = 2.868, p = 0.096 **(E)**. Fluorescence intensities of all responding TD+ neuronal cell bodies to knee compression. Pressure was increased in 50 mmHg increments every 10 s. Data represent the mean ± SEM of N = 9 cells (Cancer) from 3 animal. No between-groups comparison was done since in Sham no cells responded **(F)**. Pie chart showing the number of cells in each pressure cluster (identified in Fig. 3D) after the knee compression in FB-traced neurons in sham animals. Numbers represent cells identified in each cluster. Total numbers of cells after MCA are given in brackets **(G)**. Pie chart showing the number of cells in each pressure cluster (identified in Fig. 3D) after the knee compression in FB-traced neurons in cancer animals. Numbers represent cells identified in each cluster. Total numbers of cells after MCA are given in brackets **(H)**. Pie chart showing the number of cells in each pressure cluster (identified in Fig. 3D) after the knee compression in TD-traced neurons in cancer animals. Numbers represent cells identified in each cluster. Total numbers of cells after MCA are given in brackets. Since no TD-traced cells responded in the sham group MCA was not performed **(I)**.

### Muscle and periosteum afferents are recruited in CIBP rats

We investigated whether periosteum and muscle afferents (rather than bone afferents) are sensitised in CIBP rats by tracing muscle and periosteum (MP) afferents using injection of AAV-retrograde virus expressing tdTomato outside the tibia (Fig. 3A, 4D). We found that virtually no MP traced cells from the sham-operated rats responded to mechanical stimulation (Fig. 4E). In CIBP rats however, 20% of traced MP afferents responded to knee compression. Similarly, to total DRG cell analysis, the number of responders increased with compression intensity (Fig. 2G). This suggests that in advanced stages of the disease, cancer induces the recruitment of silent nociceptors from the tissues outside the bone rather than from the bone cavity itself. We limited the amount of virus used to ensure specificity and avoid off-target labelling in the contralateral DRG, but this mean that cell numbers were relatively low, though still considerably higher than those that can be achieved in electrophysiological studies from the same number of animals.

### Conclusions

We conclude that the increase in number of mechanically responding cells, rather than the increase in individual cell sensitivity, likely translates to the mechanical hypersensitivity observed in both CIBP rats and patients with CIBP. The mechanism appears to be activation of silent nociceptors surrounding the tumour. We also show that primary afferents, in both health and disease, respond differently according to the mechanical stimulus, suggestive of specific differential coding of pressure and proprioception by the somatosensory neurons.

## Acknowledgments

We would like to acknowledge Professor Timothy Arnett (UCL) for the use of the *µ*CT scanner, Dr Lawrence Moon (KCL) for statistical advice, Dr Douglas Lopes for manuscript assistance and Mr Alan Fieldes for professional help with MS Excel’s VBA. This work was supported by a grant from the European Union’s Horizon 2020 research and innovation programme under the Marie Sklodowska-Curie grant agreement No.642720, and by a strategic award from the Wellcome Trust (102645/Z/13/Z).

## Author Contributions

M.W.K. conceived, designed and performed all the experiments, analysed data and wrote the manuscript;

K.I.C. performed GCaMP experiments and analysed data, written R scripts for calcium imaging data analysis;

F.D. provided conceptual input on the manuscript, corrected the manuscript;

A.H.D. supervised the study, acquired funding;

K.B. provided conceptual input on the manuscript, supervised the study, co-wrote the manuscript;

S.B.M. conceived, designed, and supervised the study, corrected the manuscript, acquired funding.

## Materials and Methods

### Contact for reagent and resource sharing

Further information and requests for resources and reagents should be directed to and will be fulfilled by Stephen McMahon or Mateusz Kucharczyk.

## EXPERIMENTAL PROCEDURES

### Cell lines

Syngeneic rat mammary gland adenocarcinoma cells (MRMT-1, Riken cell bank, Tsukuba, Japan) isolated from female Sprague-Dawley rats, were cultured in RPMI-1640 medium (Invitrogen, Paisley, UK) supplemented with 10% FBS, 1% L-glutamine and 2% penicillin/streptomycin (Invitrogen, Paisley, UK). Cells were incubated at 5% CO2 in a humidity-controlled environment (37 °C, 5% CO_2_; Forma Scientific).

### Animals

Male Sprague-Dawley rats (UCL Biological Services, London, UK or Charles-River, UK) were used for experiments. Animals were group housed on a 12:12-hour light–dark cycle. Food and water were available *ad libitum*. Animal house conditions were strictly controlled, maintaining stable levels of humidity (40-50%) and temperature (22±2°C). All procedures described were approved by the Home Office and adhered to the Animals (Scientific Procedures) Act 1986. Every effort was made to reduce animal suffering and the number of animals used in accordance with IASP ethical guidelines (Zimmermann, 1983).

### Preparation of cancer cells

On the day of surgery, MRMT-1 cells were released by brief exposure to 0.1% w/v trypsin-ethylenediaminetetraacetic acid (EDTA) and collected by centrifugation in medium for 5 min at 1000 rpm. The pellet was washed with Hanks’ balanced salt solution (HBSS) without calcium, magnesium or phenol red (Invitrogen, Paisley, UK) and centrifuged for 5 min at 1000 rpm. MRMT1 cells were suspended in HBSS to a final concentration of 300,000 cells/ml and kept on ice until use. Only live cells were counted with the aid of Trypan Blue (Sigma) staining. Cell viability after incubation on ice was checked after surgery, and no more than 5-10% of cells were found dead after 4 h of ice-storage.

### Cancer-induced bone pain model

Sprague-Dawley rats weighing 120-140 g (for late-stage CIBP, 14 days post-surgery) or 180-200 g (for early-stage CIBP, 7 days post-surgery), following complete induction of anaesthesia with isoflurane (induction 5%, maintenance 1.5-2%) in 2 l/min O_2_ and subcutaneous perioperative meloxicam injection (50 μl 2 mg/kg, Metacam^®^, Boehringer Ingelheim, Berkshire, UK), were subjected to the surgical procedure of cancer cell implantation into the right tibiae(Medhurst et al., 2002). Briefly, in aseptic conditions, a small incision was made on a shaved and disinfected area of the tibia’s anterior-medial surface. The tibia was carefully exposed with minimal damage to the surrounding tissue. Using a 0.7 mm dental drill, a hole was made in the bone through which a thin polyethylene tube (I.D. 0.28 mm, O.D. 0.61 mm; Intramedic, Becton Dickinson and Co., Sparks, MD, USA) was inserted 1-1.5 cm into the intramedually cavity. Using a Hamilton syringe, either 3 × 10^3^ MRMT-1 carcinoma cells in 10 μl HBSS or 10 μl HBSS alone (Sham) was injected into the cavity. The tubing was removed and the hole plugged with bone restorative material (IRM, Dentsply, Surrey, UK). The wound was irrigated with saline and closed with Vicryl 4-0 absorbable sutures and wound glue (VetaBond 3M, UK). The animals were placed in a thermoregulated recovery box until fully awake.

### Tracing of tibial afferents

In order to study neurons innervating tibiae, rats received a single injection of a neuronal tracer into the tibial cavity a week before cancer cells implantation. An analogous surgical procedure to cancer cell implantation was used. Cancer cell injection was through the same hole in the bone to prevent further damage. Rats were injected with 5 μl of 4% Fast Blue (Polysciences Inc., Germany) solution in saline.

### Administration of tracers and calcium indicators

Male Sprague-Dawley rats weighting 60-70 g, following complete induction of anaesthesia with isoflurane (induction 5%, maintenance 1.5-2%) in O_2_ (2 l/min) were maintained at around 37°C using a homeothermic heating mat and 50 μl of Meloxicam (2 mg/kg, Metacam^®^, Boehringer Ingelheim, Berkshire, UK) was subcutaneously administered for post-operative pain management. Animals were fixed in a stereotaxic apparatus (Kopf, Germany), their lumbar region was clamped and spinal T12-L1 intervertebral space was exposed by bending the lumbar region rostrally providing easy access to the underlaying dura without the need for laminectomy. A small puncture in the dura was made and a thin catheter of 0.2 mm diameter (Braintree Scientific) was inserted in the caudal direction. 10 μl of AAV9.CAG.GCaMP6s.WPRE.SV40 (a gift from Douglas Kim & GENIE Project via Addgene viral prep #100844-AAV9, US) was infused into the intrathecal space at 1.2 μl/min (titer ≥ 1×10^13^ vg/ml). Due to the length of the inserted cannula, the infusion was close to L4 DRG. The catheter was left in place for 2 minutes before slow withdrawal. The incision was closed with wound clamps and postsurgical glue (Vetabond, 3M, UK). After 7 days of recovery, the left tibia was injected with 5 μl of 4% Fast Blue neuronal tracer (Polysciences Inc., Germany) as described above, allowing tracing of bone afferents. The muscle layer adjacent to Fast Blue injected bone was injected at the same time with 5 μl of pAAV.CAG.tdTomato (titer 7×10^12^ vg/ml, a gift from Edward Boyden via Addgene viral prep #59462-AAVrg, US) to allow tracing of muscle and periosteum afferents. After a 7-day recovery period, animals were randomly divided into two groups receiving either cancer cells or sham HBSS buffer treatment into the left tibia. Two weeks after cancer implantation animals were subjected to terminal *in vivo* calcium imaging. Throughout the whole period, body mass was carefully monitored and animals steadily gained weight resulting in 250-280 g at the day of imaging.

### *In vivo* calcium imaging of sensory neurons

Rats were anaesthetised using urethane (12.5% w/v in saline, Sigma, UK). Starting with an initial dose of 0.5 ml given i.p., subsequent doses were given at approximately 10-15-minute intervals, depending on hind limb reflex activity, until surgical depth was achieved. The core body temperature was maintained close to 37°C using a homeothermic heating mat with a rectal probe (Harvard Apparatus). Tracheotomy was performed to secure steady breathing. An incision was made to the skin on the back and the muscle overlying the L3, L4 and L5 vertebral segment was removed. The bone around either the L3 or L4 DRG was carefully removed and the underlying epineurium and dura mater over the DRG were washed and moistened with normal saline. The position of the animal’s body was varied between prone and lateral recumbent to orient the DRG in a more horizontal plane. The exposure was then stabilised at the neighbouring vertebrae using spinal clamps (Precision Systems and Instrumentation) attached to a custom-made imaging stage. The exposed cord and DRG were covered with silicone elastomer (World Precision Instruments, Ltd) to avoid drying and to maintain a physiological environment. The rat was then placed under the Eclipse Ni-E FN upright confocal/multiphoton microscope (Nikon) and the microscope stage was variably diagonally orientated to optimise focus on the DRG. The ambient temperature during imaging was kept at 32°C throughout. All images were acquired using a 10X dry objective. To obtain confocal images a 488 nm Argon ion laser line was used. GCaMP signal was collected at 500-550 nm. Time series recordings were taken with an in-plane resolution of 512 x 512 pixels and a fully open pinhole for video-rate acquisition. Image acquisition varied between 2-4 Hz depending on the experimental requirements and signal strength. At the end of the experiment, rats were sacrificed by clamping the trachea tube, left for 1h for the DRG to fill up with calcium for maximum signal control.

### Activation of sensory neurons for GCaMP *in vivo* imaging

Throughout the experiment, care was taken to provide sufficient breaks between stimuli (usually 3-5 min) in order for the tissue to equilibrate back to its baseline state. Mechanical stimulation consisted of brushing the ipsilateral calf, stretching the leg and pressure application to the leg. Brushing was performed for 10 s to the shaven surface of the cancer-bearing calf. The leg was stretched rostro-caudally, along the body axis, by 5 gentle pulling and pushing cycles. Finally, incremental pressure (50 mmHg increments every 10 s, in the range of 0-400 mmHg) was applied to the leg using a neonatal cuff connected to a manometer and air pump (air-filled 20 ml syringe). The cuff was consecutively positioned in 3 places: knee-tibial head, calf, and calf-ankle.

### Calcium imaging data analysis

Drift in time-lapse recordings was corrected using NIS Elements AR 4.30.01 (Nikon, align application). Further image processing was performed using Fiji/ImageJ Version 1.52h, and graphing and statistical analysis was undertaken with a combination of Microsoft Office Excel 2013, IBM SPSS Statistics 25 package and RStudio 0.99.893. In order to generate traces of calcium signals from time-lapse images, regions of interest (ROIs) surrounding cell bodies were chosen using a free hand selection tool in Fiji. ROIs were chosen with minimal overlap to ensure less interference from surrounding somata. A region of background was selected, and its signal subtracted from each ROI. To generate normalised data, a baseline period of fluorescence was recorded for each ROI and changes from this baseline fluorescence were calculated as ΔF/F and expressed in percentages (Chisholm et al., 2018). Implemented here are stringent criteria, where an average signal reaching 70% above baseline fluorescence plus 4 standard deviations were qualified as a response. Percentage of responders was quantified in a binary fashion within all selected ROIs. The fluorescence intensity and size analysis was performed only for responders. Non-responding cells were not analysed for their fluorescence intensity levels, as it would artificially introduce biased zero values for these cells which were either non-responding for the particular modality, or were outside of the stimulated receptive field. Thus, only those cells were analysed for intensity and size, which responded at least once to the given stimulus modality (i.e. knee compression 0-400 mmHg).

### Markov Cluster Analysis

Markov Cluster Analysis was utilised to cluster hundreds of neurons responding to the predefined stimuli (Enright, Van Dongen, & Ouzounis, 2002). BioLayout Express (under GNU Public License, Kajeka Ltd, UK) was used to run the analysis (Theocharidis, van Dongen, Enright, & Freeman, 2009). The data derived from the range of neuronal responses (ΔF/F values averaged for each stimulus length i.e. all frames from the entire duration of the 50 mmHg compression) to the defined stimuli, originating from all responders across the time. Constructed csv files with all responders’ values were used to construct network graphs (see step 1 in Fig. S3A). Initially, the similarity between individual cell responses was determined by the Pearson correlation. Pairwise Pearson correlation coefficients were calculated for every cell-set after defined sensory stimuli and correlation coefficients above a predefined threshold (R>0.9) were used to draw edges between cells (nodes) in the construction of network graphs. The nodes over the preselected value were removed from the graph (see step 2 in Fig. S3A). Several trials with different R-values were tested and the value of 0.9 was selected as the best ‘trade-off’ between overpopulated graphs with overwhelming number of clusters and no biological meanings (R<0.85) and exclusion or too many cells from the analysis (R>0.9). Next, the generated non-weighted graphs were clustered with the MCA with the following parameters: pre-inflation = 1.8, inflation = 1.8, scheme = 3, minimal number of clusters = 5. The most restrictive parameter – inflation (defines granularity of the clustering) was chosen experimentally to most tightly represent clean clusters without losing too many cells from the analysis (see step 3 in Fig. S3A). Post-hoc analysis of all visualised clusters allowed for user-defined merging of clusters with biologically-relevant similarity. This last part was supervised (see step 4 in Fig. S3A). Final core data was exported with cluster’s tag for each cell.

### Principal Component Analysis

A csv file containing final core data after MCA was used in Principal Component Analysis (PCA) to demonstrate that the classical analysis of variances is unable to detect different patterns in the longitudinal datasets (i.e. fluorescence changes to the pressure ramp across hundreds of cells). PCA was run in R (RStudio, Version 1.1.419) and analysis and visualisation were performed utilising the following packages: factoextra, corrplot, FactoMineR, ggfortify, cluster.

### Static Weight Bearing

Behaviour was assessed 2-4 hours before surgery (day 0) and at 2, 7 and 14 days following cancer cell injection. Testing was preceded by a 30 min acclimatisation period. Rooms conditions used for behavioural testing were strictly controlled, maintaining stable levels of humidity (40-50%) and temperature (22±2°C). Weight bearing was assessed using a tester (Linton Instrumentation, Norfolk, UK) in which rats were placed in a plexiglass enclosure where each hindpaw rested on a separate weighing plate. After a few minutes of habituation, the force exerted by each hind paw was measured 5 times with a 10-20 s gap between measurements. Measurements from each paw separately were averaged, and results were transformed to give the percentage of weight borne on each side to the total rear legs bearing (taken as 100%).

### Micro-computed tomography of cancer-bearing legs

Rats were sacrificed by overdose of isoflurane (5% vol/vol) and transcardially perfused with 250 ml of cold phosphate buffer saline solution (PBS, pH=7.5, Invitrogen, Paisley, UK) followed by 4% paraformaldehyde solution in 0.1 M phosphate buffer (250 mL, pH=7.5, Sigma, UK). Bones were stored frozen in -20°C until analysis. Rat tibiae, cleared of excess muscle and soft tissue, were placed into a micro-computed tomography scanner (μCT, Skyscan1172) with Hamamatsu 10 Mp camera. Recording parameters were set as follows: source voltage at 40 kV, source current at 250 μA, rotation step at 0.600 deg, with 2 frames averaging and 0.5 mm aluminium filter. For reconstruction NRecon software (version: 1.6.10.4) was used. In total, over 500, 34 μm thick virtual slices were collected per bone. Because the reference point for rat’s tibia for micro-CT analysis was not previously described, we established an anatomically relevant reference point, which was not affected by cancer, from which a region of interest was chosen for further analysis. Reference point for bone mineral density analysis (BMD) was defined as the internal tip of the intercondylar area, which was consistently located at 5 mm from the centre of the cancer growth zone. For BMD, the cancer growth zone encompassing space between 3 to 7 mm caudally from the reference point was quantified. A total of 119 scanned planes, each with a thickness of 34 μm, was analysed (see figure S1A for more details). Comparison to two known density standards allowed us to quantify BMD values in mg.cm^-3^ of both trabecular and cortical bone (utilising dataViewer software). Representative visualisations were prepared with Fiji with 3D viewer plugin.

### Immunohistochemistry

At the end of each GCaMP imaging experiment L1-L5 ipsi/contra DRG were collected and post-fixed overnight in 4% paraformaldehyde (PFA) at 4°C followed by cryoprotection in 30% sucrose (with 0.02% sodium azide) for 24 hours. After cryoprotection all DRG were embedded in Optimal Cutting Temperature (Tissue-Tek) and stored at -80°C for further analysis. The DRG embedded in OTC mould were cryo-sectioned (Bright Instruments, UK) to 10 μm thick slices collected on Menzel-Gläser Superfrost Plus Slides (25×75×1.0 mm) and stored in -20°C freezer until staining. Once dried (45°C for 2 hours) and briefly washed with 50% ethanol, sections were outlined with a hydrophobic marker (PAP pen, Japan), rehydrated and blocked with 10% donkey serum in washing solution (0.01% NaN_3_, 0.3% Triton X-100 in PBS, pH=7.5) for two hours prior to overnight incubation at room temperature with primary antibodies against GFP (to visualise GCaMP6s; chicken, 1:1000, ab13970, Abcam, UK), CGRP (marker of small peptidergic neurons; mouse, 1:1000, ab81887, Abcam, UK), IB4 (conjugated to Alexa Fluor 647; marker of small, non-peptidergic fibres; 1:250, I32450, Molecular Probes, UK), TrkA (NGF receptor; rabbit, 1:400, Abcam, ab8871, UK), Piezo2 (Piezo-type mechanosensitive ion channel component 2; rabbit, 1:200, NBP1-78624SS, NovusBio, UK). Slides were then incubated with the appropriate fluorophore-conjugated secondary antibodies (Goat anti-Chicken, Alexa Fluor 488, A11039, Invitrogen, Eugene, OR, US; Goat anti-Mouse, AlexaFluor 647, A31571, Invitrogen, Eugene, OR, US; Goat anti-Rabbit, Alexa Fluor 568, A10042, Invitrogen, Eugene, OR, US; all used at 1:1000 dilution) for 2 hours at room temperature. Slides were coverslipped using media (Fluoromount-G without DAPI, eBioscience, UK) and stored in darkness at 4°C until imaging. Samples were imaged with an LSM 710 laser-scanning confocal microscope (Zeiss) using 10x (0.3 NA) and 20 x (0.8 NA) dry objectives and analysed with Fiji Win 64.

## QUANTIFICATION AND STATISTICAL ANALYSIS

Statistical analyses were performed using SPSS v25 (IBM, Armonk, NY). All data plotted represent mean ± SEM. Detailed description of the number of samples analysed and their meaning, together with values obtained from statistical tests can be found in each figure legend. Main effects from ANOVAs are expressed as an F-statistic and P value within brackets. Statistical differences in the neuronal responses from the GCaMP experiments, were determined using a 2-way repeated-measures analysis of variance (RM-ANOVA), where applicable with Bonferroni post hoc test. Kruskal–Wallis one-way analysis of variance test (K-W, one-way ANOVA) was used to analyse behavioural data for weight bearing and body mass between Sham and CIBP groups. Unpaired t-test was used to compare number of responders in each cluster between sham and CIBP groups. One-way ANOVA with Tukey post-hoc performed in GraphPad Prism, was used to analyse data for BMD.

## Supplementary data

**Figure S1.**
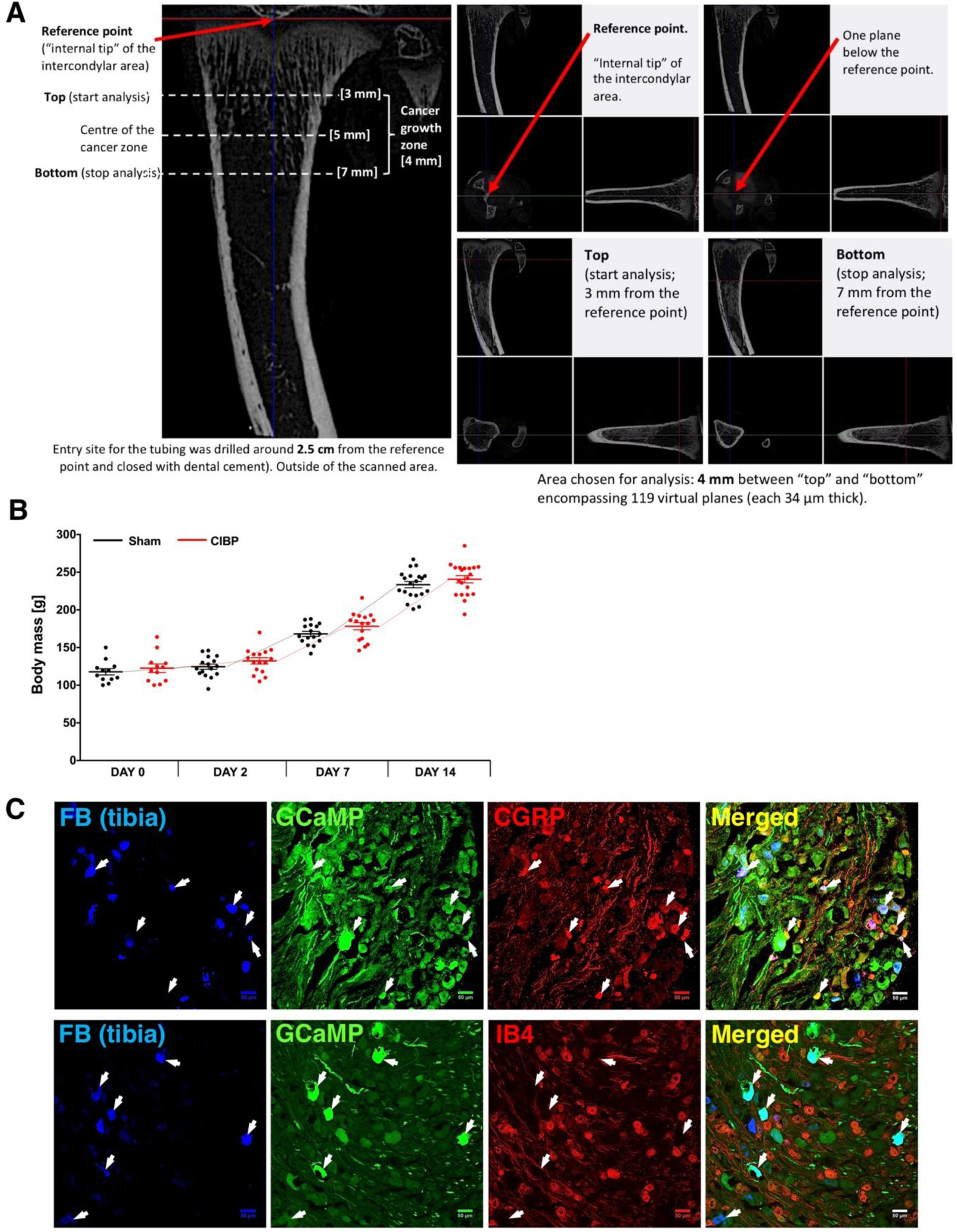
Related to Figure 1. The impact of cancer progression on bone innervation. Example micro-computer tomography reconstructions of rat tibiae. Figure explaining finding reference point for bone mineral density quantification in rat’s tibia (See methods) **(A)**. Body mass gain in rats after cancer implantation to tibia. Within a timepoint, each dot represents a single animal (n = 13-20 per group). Data represent the mean ± SEM. Kruskal-Wallis H for independent samples (Cancer vs. Sham): day 0: χ^2^(1) = 0.188, p = 0.665, day 2: χ^2^(1) = 1.503, p = 0.220, day 7: χ^2^(1) = 2.945, p = 0.086, day 14: χ^2^(1) = 1.087, p=0.297 **(B)**. Representative images of lumbar 3 (L3) DRG expressing GCaMP immunostained for peptidergic neurons marker – CGRP (top panel) and for non-peptidergic neurons marker – IB4 (bottom panel) in Fast Blue (FB) traced tibial afferents. Arrows indicate FB neurons. Scale bars, 100 *µ*m **(C)**. See also Movie S1.

**Figure S2.**
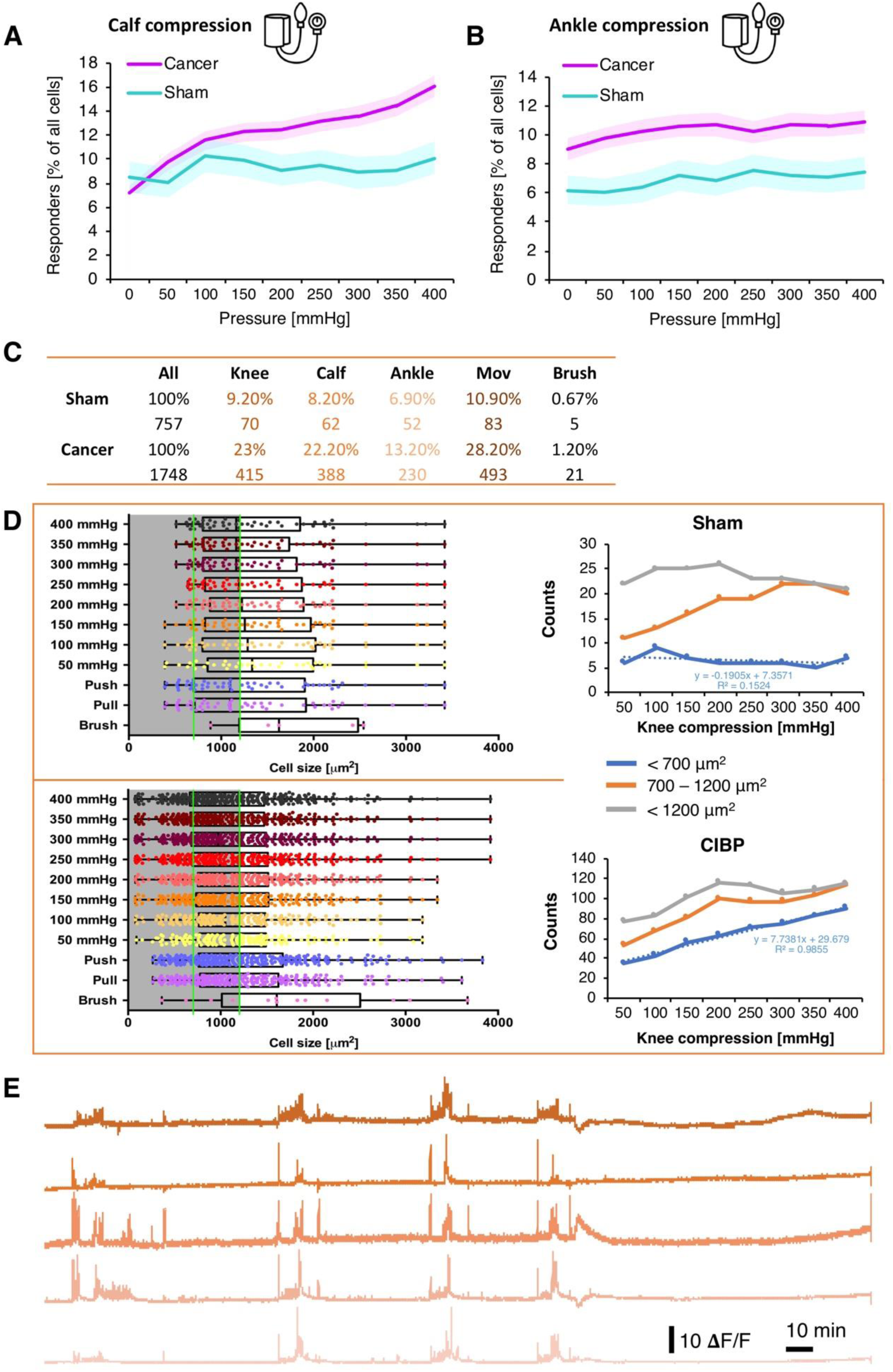
Related to Figure 2. DRG afferents encode mechanical pressure stimuli in a graded fashion. Percentage of responders during calf compression in Sham and Cancer animals. Pressure in the neonatal cuff overlaying calf was increased in 50 mmHg steps every 10 s (see methods for more). The denominator for the total number of cells was established at the end of the experiment as shown in Fig. 2C. Data represent the mean ± SEM (shaded areas) of n = 496 cells (sham) from 6 animals, and n = 1748 cells from 10 animals (Cancer). RM-ANOVA: F_1, 2242_ = 4.664, p = 0.031 (vs. sham) **(A)**. Percentage of responders during ankle compression in Sham and Cancer animals. Pressure in the neonatal cuff overlaying ankle joint was increased in 50 mmHg steps every 10 s (see methods for more). The denominator for the total number of cells was established at the end of the experiment as shown in Fig. 2C. Data represent the mean ± SEM (shaded areas) of n = 595 cells (sham) from 7 animals, and n = 1595 cells from 9 animals (Cancer). RM-ANOVA: F_1, 2188_ = 6.967, p = 0.008 (vs. sham) **(B)**. Table representing numbers of responders after each stimulus in both analysed groups. Note that the compression of the tumour-bearing area (‘knee’) results in the highest number of responding cells **(C)**. Left panels depict cell size distributions in sham (top) and CIBP (bottom) of all responders to different stimuli. Green lines highlight cell size separators used: <700 *µ*m^2^ (small cells), >700<1200 *µ*m^2^ (medium cells). Right panels show a summary count of responders to the increased knee compression within each cell size range. Sham (top), CIBP (bottom). Note that the number of small-diameter responders increase linearly with the pressure surge **(D)**. Example of 5 individual fluorescence traces (5 selected ROIs) across the whole imaging session **(E)**. See also Movie S2.

**Figure S3.**
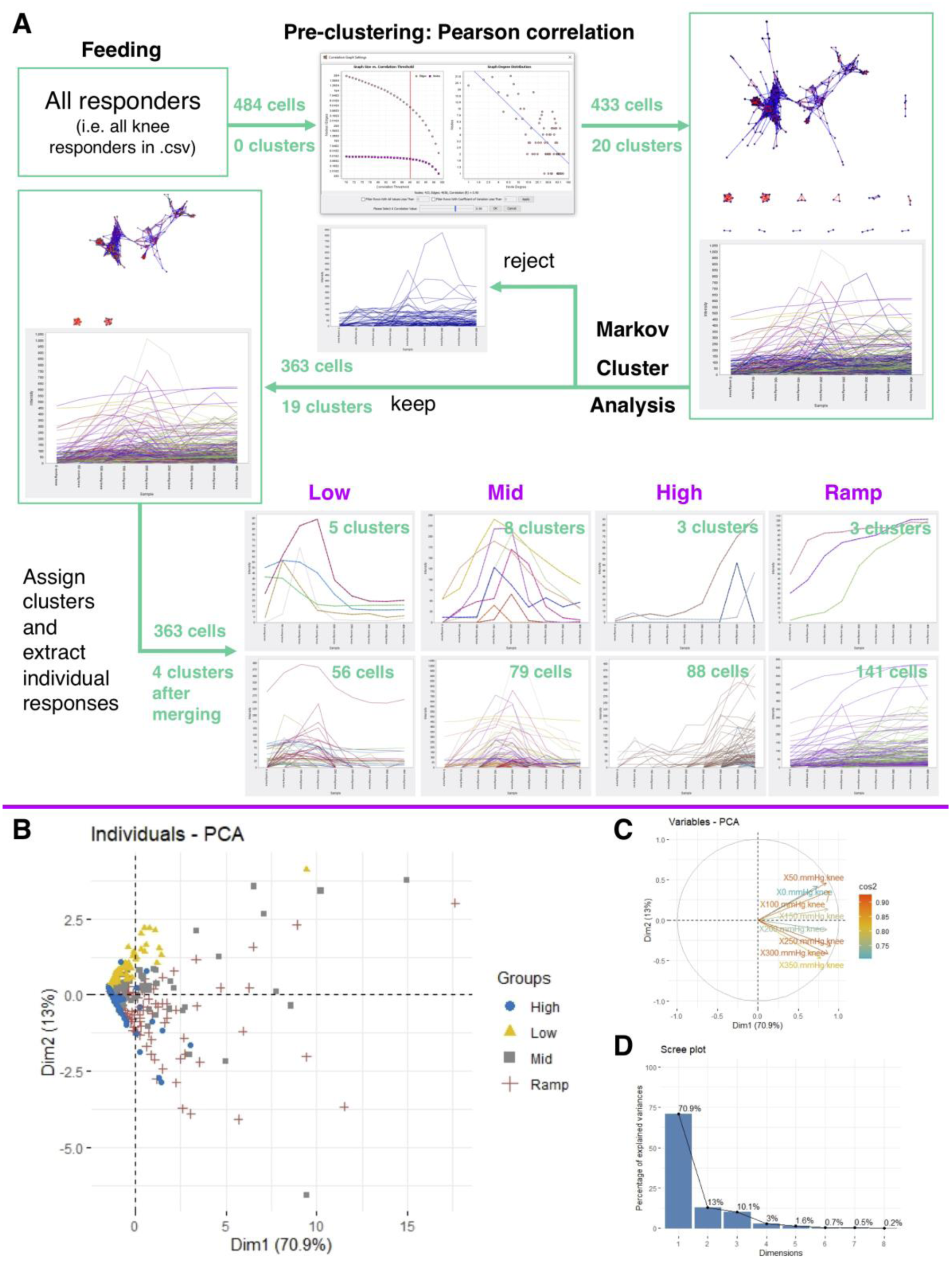
Related to Figure 3. Leg compression and position are differentially coded by DRG sensory neurons. A pipeline of the largely unsupervised Markov Clustering Analysis (MCA) of primary afferents responses. See the full description in the Material and Methods section **(A)**. Principal component analysis (PCA) was used to compare neuronal responses in pre-defined clusters. 363 knee responders fluorescence values after MCA clustering in (A) each with a cluster’s tag were used for PCA. Note that only the two most distinctive clusters (‘low’ vs. ‘high’) could be revealed with this approach (blue and yellow groups have opposite directions of the eigenvectors). The longitudinal nature of the gathered data (pressure ramp) requires advanced clustering **(B)**. Eigenvectors in the performed PCA showing the directions of the variables **(C)**. Percentage of the explained variability by each principal component after the PCA performed on the fluorescence intensity values of the knee responders **(D)**.

**Figure S4.**
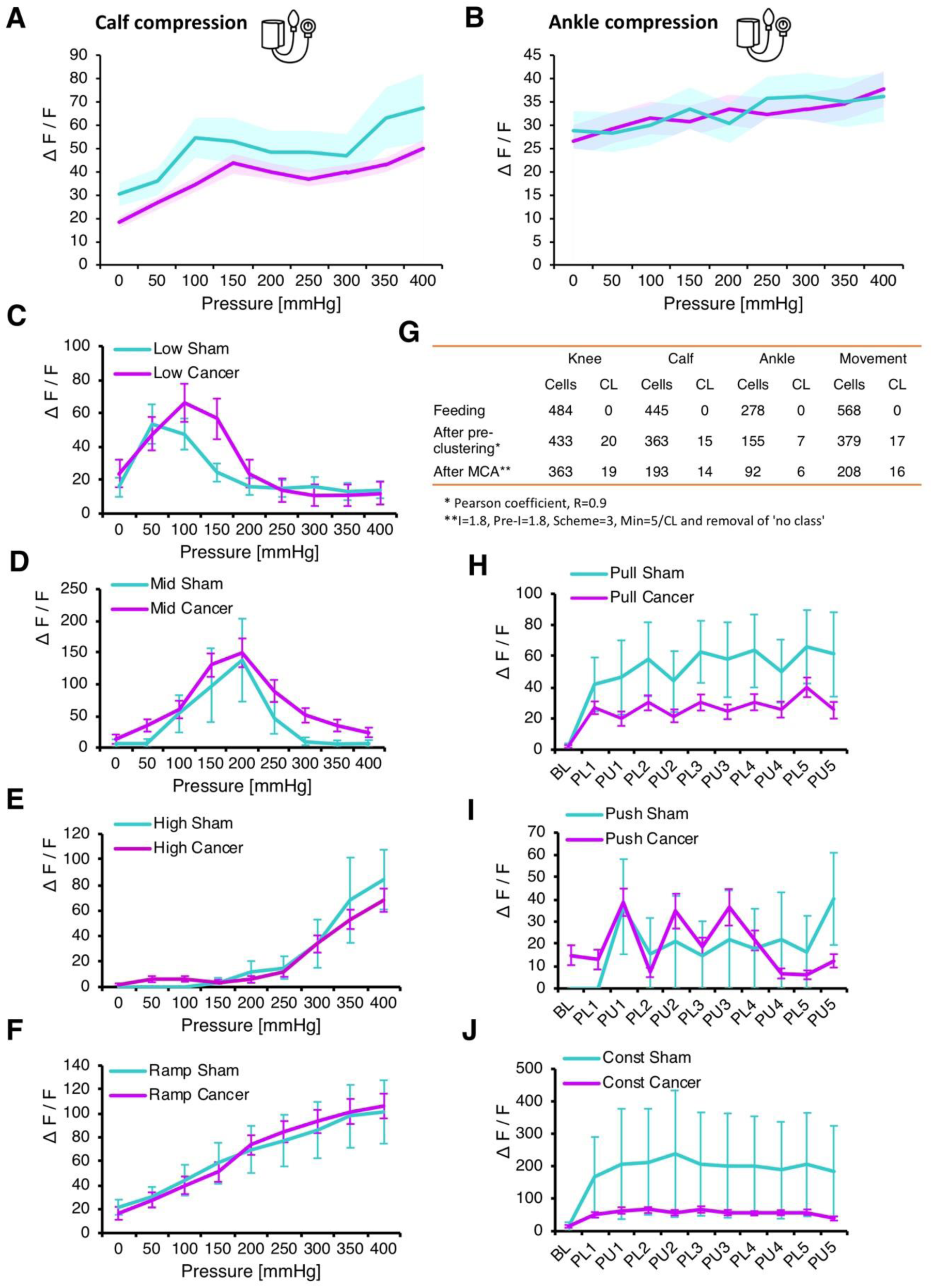
Related to Figure 3. Leg compression and position are differentially coded by DRG sensory neurons. Normalised fluorescence intensities of all responding neuronal cell bodies from L3 and L4 DRG to calf compression. Pressure was increased in 50 mmHg increments every 10 s. Data represent the mean ± SEM (shaded areas). N = 62 cells (sham) from 6 animals, and n = 388 cells from 10 animals (Cancer). RM-ANOVA: F_1, 448_ = 3.334, p = 0.069 (vs. sham) **(A)**. Normalised fluorescence intensities of all responding neuronal cell bodies from L3 and L4 DRG to ankle compression. Pressure was increased in 50 mmHg increments every 10 s. Data represent the mean ± SEM (shaded areas). N = 52 cells (sham) from 7 animals, and n = 230 cells from 9 animals (Cancer). RM-ANOVA: F_1, 280_ = 0.005, p = 0.942 (vs. sham) **(B)**. ‘Low’ cluster of neuronal responses to the knee compression revealed by Markov Cluster Analysis with respect to the treatment group. Data represent the mean ± SEM (shaded areas). N = 16 cells (sham) from 3 animals, and n = 40 cells from 10 animals (Cancer). RM-ANOVA: F_1, 54_ = 0.189, p = 0.665 (vs. sham) **(C)**. ‘Mid’ cluster of neuronal responses to the knee compression revealed by Markov Cluster Analysis with respect to the treatment group. Data represent the mean ± SEM (shaded areas). N = 5 cells (sham) from 4 animals, and n = 74 cells from 10 animals (Cancer). RM-ANOVA: F_1, 77_ = 0.303, p = 0.581 (vs. sham) **(D)**. ‘High’ cluster of neuronal responses to the knee compression revealed by Markov Cluster Analysis with respect to the treatment group. Data represent the mean ± SEM (shaded areas). N = 11 cells (sham) from 2 animals, and n = 77 cells from 10 animals (Cancer). RM-ANOVA: F_1, 86_ = 0.089, p = 0.766 (vs. sham) **(E)**. ‘Ramp’ cluster of neuronal responses to the knee compression revealed by Markov Cluster Analysis with respect to the treatment group. Data represent the mean ± SEM (shaded areas). N = 27 cells (sham) from 3 animals, and n = 113 cells from 10 animals (Cancer). RM-ANOVA: F_1, 138_ = 0.003, p = 0.959 (vs. sham) **(F)**. Summary table of all responders after each step of Markov Cluster Analysis (MCA) performed on with respect to different stimulus (See Fig. S3A). CL reflects number of clusters present at each step of the analysis before final manual merging **(G)**. ‘Pull’ cluster of neuronal responses to the leg movement along the body axis revealed by Markov Cluster Analysis with respect to the treatment group. Data represent the mean ± SEM (shaded areas). N = 17 cells (sham) from 5 animals, and n = 102 cells from 7 animals (Cancer). RM-ANOVA: F_1, 117_ = 2.278, p = 0.134 (vs. sham) **(H)**. ‘Push’ cluster of neuronal responses to the leg movement along the body axis revealed by Markov Cluster Analysis with respect to the treatment group. Data represent the mean ± SEM (shaded areas). N = 4 cells (sham) from 3 animals, and n = 33 cells from 6 animals (Cancer). RM-ANOVA: F_1, 35_ = 0.001, p = 0.970 (vs. sham) **(I)**. ‘Const’ cluster of neuronal responses to the leg movement along the body axis revealed by Markov Cluster Analysis with respect to the treatment group. Data represent the mean ± SEM (shaded areas). N = 2 cells (sham) from 2 animals, and n = 25 cells from 5 animals (Cancer). Statistical comparison not performed due to the low cell numbers in sham group **(J)**.

**Figure S5.**
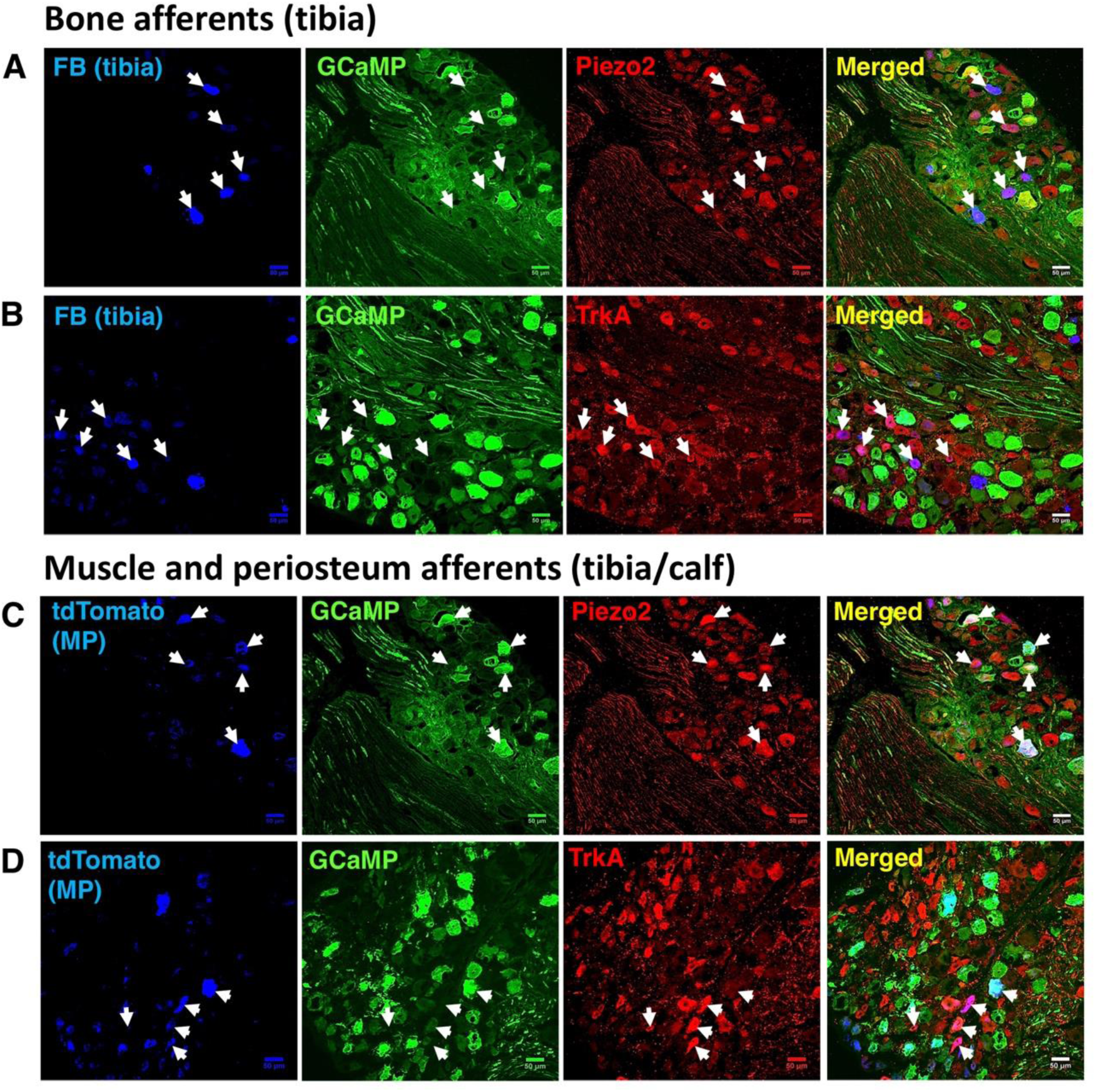
Related to Figure 4. Intra- and peri-tibial afferent function in health and bone cancer. Representative images of Fast Blue (FB) traced tibial afferents in L3 DRG collected after the GCaMP experiments. Sliced DRG were immunostained for Piezo2 **(A)** and TrkA **(B)**. Representative images of AAVrg/tdTomato (tdTomato) traced muscle and periosteum (MP) afferents in L3 DRG collected after the GCaMP experiments. Sliced DRG were immunostained for Piezo2 **(C)** and TrkA **(D)**. Scale bars: 50 *µ*m.

